# *Saccharomyces cerevisiae* adapted to grow in the presence of low-dose rapamycin exhibit altered amino acid metabolism

**DOI:** 10.1101/368266

**Authors:** Duygu Dikicioglu, Elif Dereli Eke, Serpil Eraslan, Stephen G Oliver, Betul Kirdar

## Abstract

Rapamycin is a potent inhibitor of the highly conserved TOR kinase, the nutrient-sensitive controller of growth and aging. It has been utilised as a chemotherapeutic agent due to its anti-proliferative properties and as an immunosuppressive drug, and is also known to extend lifespan in a range of eukaryotes from yeast to mammals. However, the mechanisms through which eukaryotic cells adapt to sustained exposure to rapamycin have not yet been thoroughly investigated. Here, *S. cerevisiae* response to long-term rapamycin exposure was investigated by identifying the physiological, transcriptomic and metabolic differences observed for yeast populations inoculated into low-dose rapamycin-containing environment. The effect of oxygen availability and acidity of extracellular environment on this response was further deliberated by controlling or monitoring the dissolved oxygen level and pH of the culture. Yeast populations grown in the presence of rapamycin reached higher cell densities complemented by an increase in their chronological lifespan, and these physiological adaptations were associated with a rewiring of the amino acid metabolism, particularly that of arginine. The ability to synthesise amino acids emerges as the key factor leading to the major mechanistic differences between mammalian and microbial TOR signalling pathways in relation to nutrient recognition. Furthermore, oxygen levels and extracellular acidity of the culture were observed to conjointly affect yeast populations, virtually acting as coupled physiological effectors; cells were best adapted when maximal oxygenation of the culture was maintained in slightly acidic pH, any deviation necessitated more extensive readjustment to additional stress factors.

## INTRODUCTION

The target of rapamycin (TOR) serine/threonine kinase is a highly conserved protein, which is the nutrient-sensitive central player in controlling growth and aging [1]. First discovered in the yeast, *Saccharomyces cerevisiae*, TOR is a functionally conserved protein across eukaryotes in Kingdoms Fungi, Plantae, and Animalia, from yeast to human [2]. TOR signalling pathway was shown to stimulate growth in response to increased nutrient availability [3] by suppressing stress response to further promote growth and proliferation [4].

Rapamycin is a potent inhibitor of TOR, discovered to inhibit the proliferation of the infectious yeast *Candida albicans* [5], and has thus far been utilised as a chemotherapeutic agent with anti-proliferative properties and as an immunosuppressive drug [6]. Rapamycin was reported to affect protein synthesis *via* blocking the initiation of translation and altering the phosphorylation of the translation-related proteins [7–9]. Additionally, the transcriptional changes associated with rapamycin treatment were shown to be similar to those associated with nutrient starvation [1, 9, 10]. The inhibition of TOR activity by low-dose rapamycin was shown to extend both chronological and replicative lifespan of *S*. *cerevisiae* [11], and to improve the survival of cancer-prone mice, and of mice suffering from premature aging [12–15].

Investigation of the cellular response of yeast cells to rapamycin inhibition of TOR focused primarily on monitoring the short-term effects of the treatment, where the response to a range of rapamycin concentrations (0.2 nM – 400 nM) was investigated within three hours post-treatment [16–23]. The recovery of yeast cell populations from the anti-proliferative effect of rapamycin was also investigated, and the EGO complex was shown to play an important role in the process [24]. Microbial systems are very successful in adaptation by rewiring their physiology in response to sustained exposure to mild stresses [25–27]. Stress-inducing conditions could even lead to evolved populations if the acquired advantage were also accompanied by fitness rewards [28]. However, the transcriptional and metabolic landscape of microbial populations adapted to grow in the presence of low concentrations of rapamycin, known for its growth inhibitory effects, has not yet been explored in this context, despite numerous reports on its ability to extend lifespan [2, 5, 29, 30].

Multiple cross-regulatory interactions were identified between the TOR and the Mitogen Activated Protein Kinase (MAPK) signalling pathways in the fission yeast and these cross-talk were proposed as mechanisms to elicit adaptive responses to extra- and intracellular conditions by regulating essential cellular functions [31].

Since MAPK signalling pathways, particularly those of mating and filamentous growth, were shown to be responsive to intermediary levels of oxygen availability in the baker’s yeast [32], whether TOR-induced cellular events could also to be responsive to oxygen levels poses an interesting query in light of these recent findings.

Calorie restriction-induced extension of lifespan in *S. cerevisiae*, which prefers a slightly acidic environment where cellular proliferation was shown to occur rapidly, was reported to be achieved by reducing fermentative growth and secreting fewer organic acid molecules. This allowed the maintenance of relatively high extracellular pH [33]. High extracellular acetic acid levels combined with low pH was likewise reported to be toxic and accelerate yeast mortality. Consequently, reduction in TOR signalling was reported to enhance respiratory activity and limit acetic acid production by shunting fermentative products into respiratory metabolism [34]. Contrariwise, syncing the extracellular and the intracellular pH was also proposed as a possible strategy that yeast employs to decrease the inhibitory effect of organic acids, and associated cellular stress. This would, in turn, reduce energy consumption for maintaining the optimal intracellular pH, and thus significantly increase the chronological life span in yeast [33, 35, 36]. Regardless, the interplay between extracellular pH and yeast lifespan has not been explored during the course of chemical inhibition of TOR.

In this work, we ask the following questions: (i) Is the adaptive response of yeast to sustained and long-term rapamycin exposure different from its short-term rapid response? (ii) Does this response vary as extracellular environment is more or less acidic or as oxygen levels vary? For this purpose, we identified the growth characteristics of yeast cultures grown and maintained in the presence of low doses of rapamycin. We then investigated the physiological, transcriptomic and metabolic response of yeast populations inoculated into and cultivated in low-dose rapamycin-containing environment. This analysis was conducted in cultures where the oxygen availability and the level of acidity was either controlled or monitored as set by 2 2 factorial design. The investigation of the effect of oxygen availability and the pH of the extracellular environment on rapamycin treatment was shown to be particularly important in understanding the mechanisms of adaptation to long-term rapamycin exposure.

## MATERIALS AND METHODS

### Strain and growth conditions

Homozygous *hoΔ*/*hoΔ* strain and wild type strain of *S. cerevisiae* BY4743 (*MATa*/*MAT*α *his3Δ1*/*his3Δ1 leu2Δ0*/*leu2Δ0 lys2Δ0*/+ *met15Δ0/*+ *ura3Δ0*/*ura3Δ0*) were purchased from EUROSCARF. The genetic background of the *hoΔ*/*hoΔ* strain was verified using PCR-based methods [54]. All precultures were grown overnight in rich medium (2% [w/v] D-glucose, 2% [w/v] peptone, 1% [w/v] yeast extract) at 30°C in an orbital shaker shaking at 180 rpm prior to inoculation. Defined synthetic medium [55] was used for the cultivations carried out in the fermenters.

### Determination of the working concentration of rapamycin

Cells were grown in identical shake flasks containing rich medium with 1:7 working volume to total volume ratio at final concentrations of 0.5nM, 2nM, 5nM, 10nM, 20nM, 100nM, 200nM, 300nM, 400nM, 500nM, 700nM and 1000nM of rapamycin (Sigma Cat. No: R0395) dissolved in 90/10 (v/v) ethanol/Tween-20 solution. Rapamycin was either introduced prior to inoculation, or during the exponential phase of the culture growth. The optical density (OD_600_) was monitored throughout cultivation spectrophotometrically (DU730, Beckman Coulter Inc., U.S.A.), and the final biomass concentrations were determined gravimetrically. The chronological lifespan was determined as described in [56] by daily sampling the culture after 72 h *via* serial 1:20 dilutions of ca. 2×10^5^ cells and monitoring colony growth on solid synthetic or rich medium after 3 or 2 days, respectively.

### Fermentations, sampling and determination of growth characteristics

Batch cultivations were carried out in 2L B-Braun Biostat B Plus fermenters in duplicates (1.5L working volume maintained at 30°C with the rate of agitation at 800rpm). The dissolved oxygen (dO_2_) saturation was maintained > 90% by constant air flow at 1.5 L/min in experiments where air supply into the culture was controlled (air^+^). Micro-aerated cultivation vessels were brought to dO_2_ saturation of 100% prior to inoculation and air supply was then blocked (air−). pH was either maintained at 5.5 *via* 2-point control by NaOH and HCl (pH+), or was monitored (pH-) to follow its own course. Samples were collected for endo/exo-metabolite analysis and gene expression analysis as well as for biomass determination. The samples for transcriptome and metabolite analyses were collected at OD_600_ range of 0.6–0.8. Biomass density was determined gravimetrically at the time of sampling for analytical and gene expression measurements and at stationary phase. Supernatants were stored at –20°C until exo-metabolite analysis. Cells harvested for microarray analysis were immediately frozen in liquid nitrogen and were stored at –80°C until further processed.

### RNA isolation and microarray analysis

Total RNA was isolated in QIAcube robotic workstation operating a modified mechanical lysis protocol following the RNeasy mini kit (Cat no: 74106) protocol (Quiagen U.S.A.).

RNA quantification and purity (A_260_/A_280_) evaluation was carried out using a micro-volume UV-vis spectrophotometer (NanoDrop ND-3000, Thermo Fisher Scientific Inc., U.S.A.). RNA integrity was checked in a microfluidics-based platform (Bioanalyzer 2100) using RNA6000 Nanokit (Agilent Technologies, U.S.A.).

Microarray analysis was performed as described in the GeneChip®Expression Analysis Technical Manual (relevant kits, chips, and instrumentation: Affymetrix Inc., U.S.A.). Briefly, first-strand cDNA was synthesized from *ca*.100ng of total RNA, and converted into ds DNA (GeneChip® 3’ IVT Express Kit). Biotin-labelled aRNA was synthesised, purified and fragmented with the relevant quality control measures in place prior to loading 5μg of aRNA onto 169 format Yeast 2.0 arrays. The chips were washed and stained in the GeneChip® Fluidics Station, and GeneChip® Scanner 3000 was used for image acquisition. The image files were processed and normalised with their quality assessed by dChip software [57]. MIAME [58] compliant raw and processed files can be accessed from EBI’s ArrayExpress database under the accession number E-MTAB-6628 (https://www.ebi.ac.uk/arrayexpress/).

### Metabolite analyses

Sample preparation for exo- and endo-metabolite measurements were carried out as described previously [59]. The supernatant stored at –20°C was used for the analysis of extracellular metabolites. Cells were subjected to dry ice / methanol quenching, and the intracellular metabolites were extracted by the boiling ethanol protocol. The extracts were lyophilised and stored at −20°C until further analysis.

Chromatographic separation of amino acids was performed using an Agilent 1260 Infinity UPLC (U.S.A.) with a 4.6 × 100 hydrophilic interaction chromatography (HILIC) column (Zorbax Rapid Resolution, particle size 3.5µM). The initial composition of the mobile phases were 15% (A) containing 40mM ammonium formate with 2% formic acid and 85% (B) acetonitrile, followed by a linear gradient to 85% A and 15% B in 13 minutes with a two-minute hold and four-minute re-equilibration of the column back to its initial conditions prior to the next run amounting to a total assay time of 19 minutes. The flow rate was set at 0.4µL/min and 2µL of sample was injected.

Chromatographic separation of TCA cycle metabolites was performed using an Agilent 1260 Infinity UPLC (USA) with a 4.6 × 150 Eclipse XDB column (Zorbax Rapid Resolution, particle size 5µM). The initial composition of the mobile phases were 15% (A) containing 10mM ammonium acetate adjusted to pH 8 using ammonium hydroxide and 85% (B) acetonitrile with 0.1% formic acid, followed by a linear gradient to 85% A and 15% B in 4 minutes and back to its original composition immediately after 4 minutes with a hold run for another 6 minutes for the re-equilibration of the column prior to the next run amounting to a total assay time of 10 minutes. The flow rate was 0.4 µL/min with a 2µL sample injection.

All standards employed in spectrometric analysis were obtained from Sigma-Aldrich, Inc. (U.S.A.). The optimal MS/MS MRM transitions, fragmentation patterns and the retention time behaviours of each metabolite were determined using Agilent 6430 Triple Quadrupole –MS (U.S.A.). The MS parameters were as follows: gas temperature at 300°C, gas flow rate of 10L/min, nebulizer pressure of 40 psi, and capillary current at 3750V. The ESI probe was operated at positive ion mode for amino acids and negative mode for TCA cycle metabolites. The quantitation of the samples was acquired using MassHunter Quantitative Analysis Software, version B.05.00 (Agilent Tech. Inc., U.S.A.) against calibration curves generated with standard solutions.

The glucose, glycerol, ethanol and ammonium content of the supernatant were determined enzymatically by rbiopharm Roche Yellow Line assays (Cat nos: 10716251035, 10148270035, 10176290035, and 11112732035, respectively) as described by the manufacturer.

### Data analysis

Student’s t-test was employed for statistical analysis whenever sample size ≥ 3, followed by the Benjamini-Hochberg test when sample size > 100 and by Bonferroni correction when sample size < 100 to control the False Discovery Rate. 1-way ANOVA and 2-way ANOVA were employed as necessary for datasets where samples size < 3, followed by Tukey HSD or Sidak’s test as post-hoc analysis for multiple comparisons. Significance was evaluated at the confidence level of α = 0.05. A fold change threshold of 1.5 was employed in the magnitude of gene expression in order to identify substantial changes. Hierarchical clustering of samples was carried out using Hierarchical Clustering Explorer (HCE) 3.5 [60], with Pearson correlation employed as the distance metric and average linkage employed as the linkage metric. Princeton GO tools were employed for Gene Ontology Enrichment Analysis (http://go.princeton.edu/cgi-bin/GOTermFinder) accessed on (02/2018) [61]. At least 4 replicate measurements of each population were employed in the significance testing of shake flask experiments.

## RESULTS

### Continuous exposure to low-dose rapamycin boosts culture growth and extends chronological lifespan

We initially investigated the effect of exposing yeast to rapamycin for extended periods on the growth phenotype of the cultures since rapamycin is a potent inhibitor of TOR kinase, the major regulator of growth (see ESM1 for the complete dataset). For this purpose, two different modes of rapamycin administration were evaluated: (i) cells were directly inoculated in media initially containing rapamycin; denoted as *ab initio* treatment, and (ii) rapamycin was added into the culture during exponential growth phase, i.e. induction by rapamycin (Fig. 1a). Cellular growth was evaluated using the optical density (OD) measurements.

**Fig.1.**
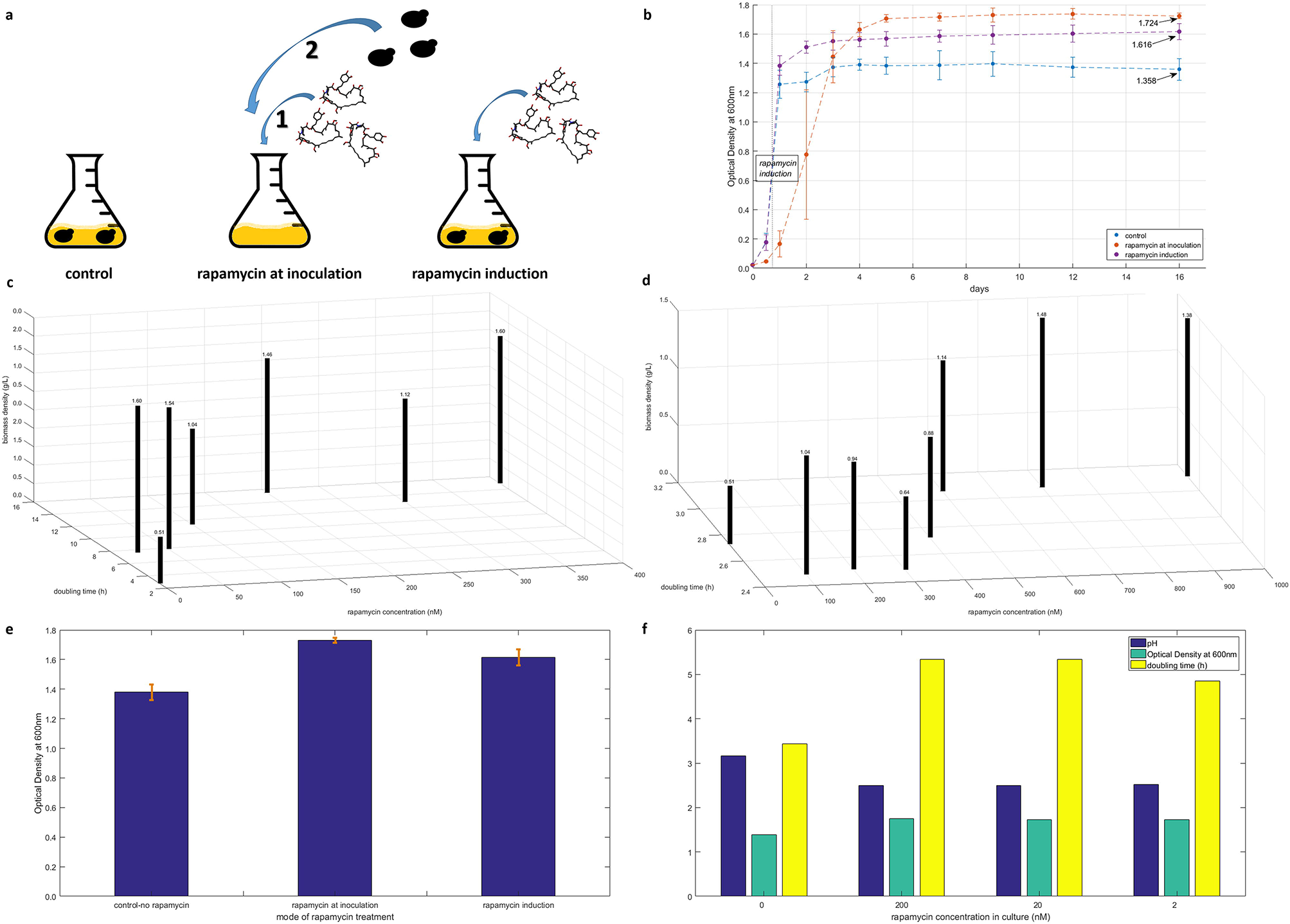
Optimisation of the working concentration and the mode of administration of rapamycin Schematic representation of the experimental setup for investigating the effects of different modes of rapamycin treatment where the cells are either introduced to rapamycin-containing medium, or a culture at steady growth was induced by rapamycin (a). Growth profiles of yeast culture under the tested conditions. The optical density (OD) of the cultures was plotted against the number of days of culturing. The OD at which rapamycin induction was done was designated by the “rapamycin induction” box. The final undiluted ODs for each mode of treatment on day 16 was designated by an arrow (b). The average biomass density of the cultures inoculated into rapamycin containing medium (c) and that of rapamycin-induced cultures (d) were plotted against the rapamycin concentration employed in the treatment and the doubling time of the culture calculated from its maximum specific growth rate. The numerical values for biomass density are indicated above each bar. The variation in the final undiluted OD values among cultures that were subjected to the same mode of treatment were highlighted in orange; despite all variation being within acceptable limits, that of cells inoculated into rapamycin containing medium were remarkably lower than that of the control and the other mode of treatment (e). The effect of lowering the working concentration of rapamycin (on the abscissa) on the final extracellular pH of the culture (column in navy), the final OD value reached (column in teal) and the doubling time during balanced growth (column in yellow) are provided in a range of 2 orders of magnitude: 2–200nM (f).

A range of rapamycin concentrations (20 – 1000nM) were tested, and *ab initio* treatment delayed exit from lag phase significantly by 133% (p-value = 7.14 × 10^−12^). The doubling time (t_D_) of these cultures was significantly higher by 4.2-fold (p-value = 8.38 × 10^−6^) than the control cultures and the rapamycin-induced cultures during the exponential phase of growth, and these effects were observed to be independent of the concentration of rapamycin used. Yeast cultures were observed to reach significantly higher cell densities in the presence of rapamycin than in control cultures regardless of the mode of administration (OD_600_ = 1.358 vs 1.667, p-value = 2.79 × 10^−5^). *Ab initio* treated yeast cultures reached even higher final cell densities than the rapamycin-induced populations despite the delayed and decelerated growth in early phases (OD_600_ = 1.616 vs 1.724, p-value = 2.35 10^−4^) (Fig. 1b). The control cultures were observed to lose viability starting from day 7, and they were completely inviable by day 12, as were the rapamycin-induced cultures. The *ab initio* treated cultures survived for a further 4-day period, and by the 16^th^ day, they have completely lost viability.

The final biomass density of the *ab initio* treated cultures did not demonstrate an increasing or a decreasing trend as the rapamycin concentration of the medium was increased from 20nM to 400nM. However, the t_D_ of the cultures were shorter, particularly for cultures where rapamycin concentration was below 100nM (Fig. 1c). In contrast, the cultures, which were induced with rapamycin, had an increasing t_D_ and final biomass density as rapamycin concentration was increased, particularly up to 500nM (Fig. 1d). The only outlier in both cases was those of cultures treated with 300nM rapamycin. The variability of the cultures under different treatment regimes revealed minor but distinct differences. The variation in replicate cultures was observed to be much less for *ab initio* treated cultures than those for control or rapamycin-induced cultures (Fig. 1e).

These observations led to further exploration of low rapamycin concentrations and inoculation into rapamycin-containing medium since both the early phase slow-growth characteristics and the final cell growth and chronological lifespan measurements associated with this system indicated a remarkably different growth phenotype for the yeast population. Reducing the concentration of rapamycin employed by a magnitude (2nM working concentration) did not change the growth characteristics except for only a marginal decrease in doubling time (Fig. 1f). Furthermore, the final extracellular pH of the culture was a unit lower than that of control samples in the presence of rapamycin, but this observation did not show a concentration-dependent trend (Fig. 1f). The reduction of the working concentration further down to 0.5nM did not yield any significant difference in final cell density (p-value = 0.38); nevertheless the working concentration was set at 2nM for the following experiments.

### Physiological response of yeast cells cultured in low-dose rapamycin-containing environment

Having established the 2nM working concentration of rapamycin and a mode of exposing the cells to the inhibitor by inoculating the cells into rapamycin-containing medium, we then designed a setup to investigate the physiological, and the cellular response of yeast cells. Rapamycin-exposed cultures were observed to reach lower extracellular pH than control cultures in our preliminary experiments, possibly by enhanced secretion of organic acids. In light of these results, the cultivations were conducted either (i) by controlling the culture pH at a slightly acidic level (pH = 5.5, pH+) lying within the widely accepted optimal range for *S. cerevisiae* growth, or (ii) by monitoring the pH profile across time without introducing any control actions (pH-). Oxygen availability was introduced as an additional factor; the cultures were either (i) controlled at full-saturation (> 90%) of dissolved oxygen (dO_2_) (air+), or (ii) brought to full saturation prior to inoculation and monitored through the course of culture growth as oxygen availability decreased (air-). Growth characteristics of these cells and extracellular concentration of substrates and by-products including alcohols, amino acids and tricarboxylic acid cycle (TCA) intermediates were measured in order to understand the physiological changes the cells undergo as a response to sustained exposure to rapamycin (ESM2).

The biomass density of the cultures measured at stationary phase were significantly lower (p-value < 2.88 × 10^−3^) for rapamycin-treated cells, except for the air-pH- cultures, where the biomass density was significantly higher than those of control cultures (p-value = 1.07 × 10^−2^), as observed in the shake-flask experiments, the closest prototype of the air-pH- cultures. The rapamycin-treated cultures were observed to reach mid-exponential phase significantly slower than untreated cultures (p-value = 5.88 × 10^−3^), and the glucose uptake of these respective cultures was significantly lower than that of the control cultures (p-value = 2.79 × 10^−3^) (Fig. 2a). This was caused by an extended lag phase, in line with the observations from shake-flask experiments, but it was not coupled with a lower specific growth rate. Furthermore, the difference in the length of the lag phase was not as pronounced as in shake flask experiments. Differences in the rate of agitation was identified as the main cause of this observation. The rate of agitation in the fermenters was nearly 4.5-fold faster than in orbital shakers, allowing superior mixing and improved oxygen transfer rates even for air-cultures. We verified this notion by conducting rapamycin treated air-pH-fermentations (as in orbital shakers) reducing the rate of agitation from 800rpm to 400rpm. A time delay of 4 hours (ca. 2 t_D_ s) was observed at 400 rpm to exit lag phase and the specific growth rate was slowed down by 36% confirming our hypothesis (see ESM2 for details).

**Fig.2.**
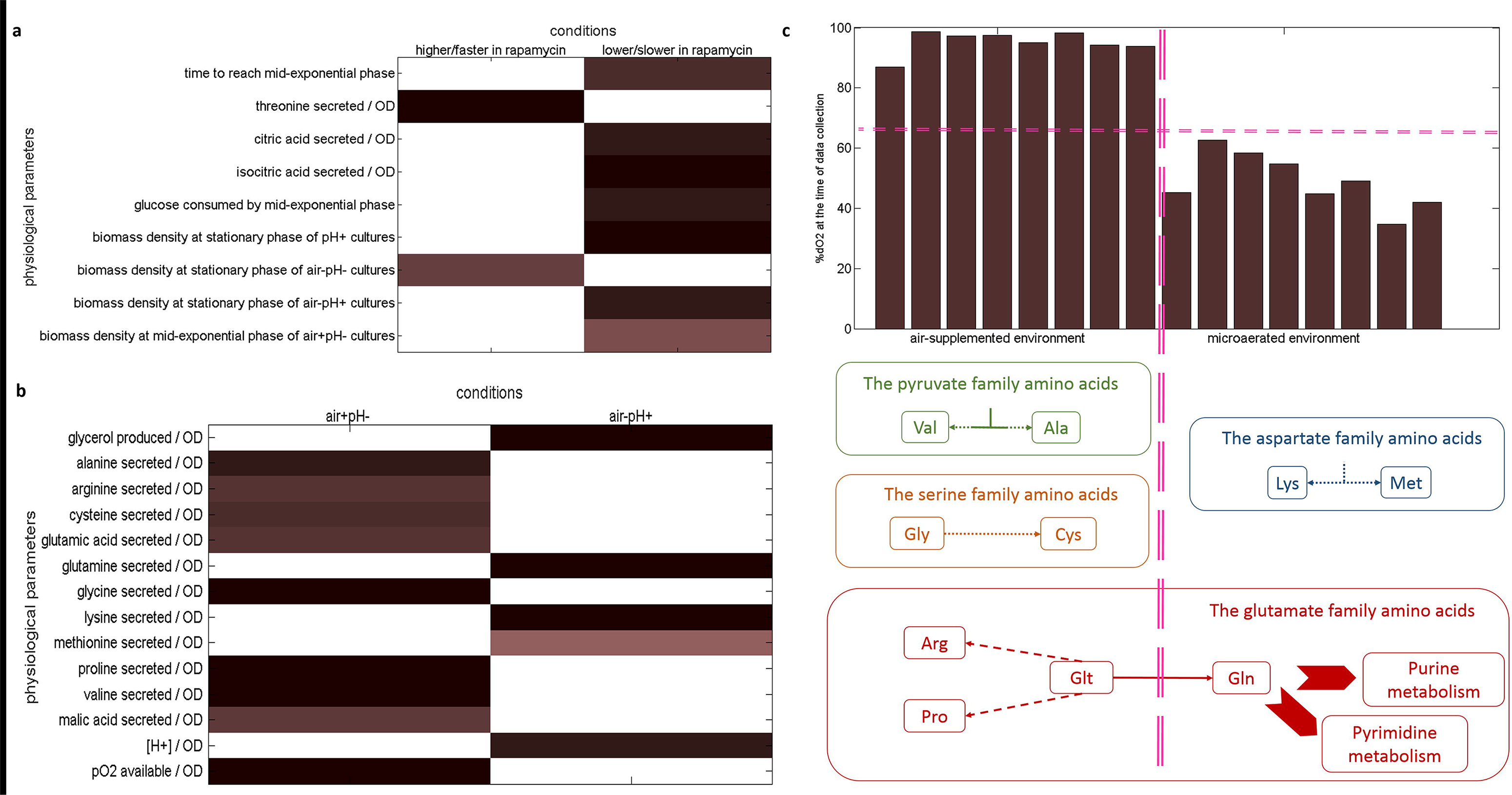
Overall analysis of the physiological response of yeast to long-term exposure to rapamycin Physiological parameters that showed a significant difference in response to rapamycin are displayed in (a). The two columns in the plot separate those responses that were higher (or faster depending on the parameter) in yeast cultures that were exposed to rapamycin, from those that were lower (or slower). Metabolite yields that showed a significant difference in air+pH- and air-pH+ cultures are displayed in (b). The two columns in the plot separate those responses that were higher yields achieved in either air+pH− or air-pH+ cultures. Colour white denotes non-significant measurement in both plots (a) and (b), and deeper the shade (towards mahogany), higher the significance of the evaluation (i.e. lower the p-value of the test statistic). Summary of the findings in (b) within the context of fermenter dO_2_ levels is displayed in (c). The amino acids whose yields changed significantly between air+pH− and air-pH+ cultures are structured into families based on their ancestor molecule (colour-coded similarly), and these responses are coupled with a plot displaying the dO_2_ availability of these cultures at the time of sampling (top display). The horizontal magenta dashed line accentuates the difference in oxygen availability at the time of sampling, and the vertical magenta dashed line separates responses for air+ and air− cultures.

The presence of rapamycin in the cultures where only dO_2_ or pH was controlled (i.e. the air+pH− and air-pH+ cultures) induced significant differences in the cell densities (p-value = 5.30 × 10^−2^ *and* 3.84 × 10^−2^ for air+pH− and air-pH+, respectively) while no such difference was observed in the rapamycin-containing air+pH+ and air-pH- cultures. This led to the investigation of metabolite yields of these cultures on cell density, in order to account for this difference. Only extracellular threonine, isocitrate and citrate yields were observed to be significantly different between the treated and the untreated cultures (p-values = 2.59 × 10^−4^, 2.32 × 10^−3^, 7.95 × 10^−4^, *respectively*) (see Fig. 2a). However, owing the differences observed in cell density between control and rapamycin cultures, the yields for a number of extracellular metabolites, majority of which were amino acids, were observed to be significantly different between air+pH− and air-pH+ cultures (p-value = 2.50 × 10^−2^) in response to differences on oxygen or pH levels (Fig. 2b). The amino acids, whose yields significantly differed between conditions, did not show any trends in terms of their properties such as their size, shape, solubility, or ionization properties of their R groups, which would substantially influence the structure and biological activity of proteins that they constitute. However, a closer inspection of these amino acids highlighted their clustering into families that were derived from a common molecule. Many of these amino acids appeared to be driven from the TCA cycle metabolites as their precursors. In light of these observations, the aeration level of the cultivations could potentially be a greater influence than the pH of the extracellular environment on the type of amino acids secreted from yeast under these conditions, since their cognate metabolic activities were centred on the TCA cycle. The following evaluations were accordingly carried out to investigate the impact of aeration level on these systems. The yields of valine and alanine from the pyruvate family, as well as of glycine and cysteine from the serine family were all significantly higher in cultures controlled at 100% dO_2_ saturation (p-value < 5 × 10^−3^), whereas the yields of lysine and methionine from the aspartate family were significantly higher at ca. 50–60% dO_2_ saturation (p-value < 2.50 × 10^−2^). The yields of the glutamate family amino acids were unpredictable. The yields of arginine, proline and glutamic acid, which are known protein synthesis precursors, were significantly higher when oxygen level was controlled at full saturation (p-value < 8 × 10^−3^). On the other hand, the yield for glutamine, a precursor for the biosynthesis of purine and pyrimidine nucleotides, was higher at low oxygen saturation (p-value = 3.03 × 10^−5^) (Fig. 2c). The preferential activation towards protein or nucleotide biosynthesis implicated by the generation of suitable amino acid pools was in line with phosphoproteomic data reporting the yeast TORC1 to be responsible for the regulation of nucleotide and amino acid biosynthesis [23]. Although the conclusions above were drawn from analyses carried out pooling treated and untreated cultures, a similar analysis conducted on untreated or treated cultures alone did not yield a similar outcome indicating that these differences that become more prominent as a response to oxygen levels was indeed caused by factoring rapamycin treatment into the equation, and was not only caused by differences in oxygen levels.

Physiology of the yeast cells were extensively modified as a response to long-term rapamycin exposure, and this response was coupled with the dissolved oxygen level and the extracellular pH of the cultures from which these data were collected. We conducted ANOVA to identify these culture-specific responses and identified that 75% of the physiological parameters under investigation (33/44) were significantly altered upon sustained exposure to rapamycin in at least either one of the four cultivation conditions. However, majority of the responses that were significantly higher (or faster as the parameter implies) in yeast cultures that were exposed to rapamycin (p-value < 5.00 × 10^−2^) (Fig. 3a), and those that were lower (or slower) (p-value < 3.00 × 10^−2^) (Fig. 3b) were exclusive, except for only three parameters; extracellular methionine, threonine and ethanol levels at mid-exponential phase. The most number of significant changes were observed around cultures where dO_2_ and pH were both controlled (air+pH+; 17 parameters) and around those where neither of these were controlled (air-pH-; 15 parameters), suggesting a coupled extracellular oxygen-acidity effect on the yeast response to sustained exposure to rapamycin.

**Fig.3.**
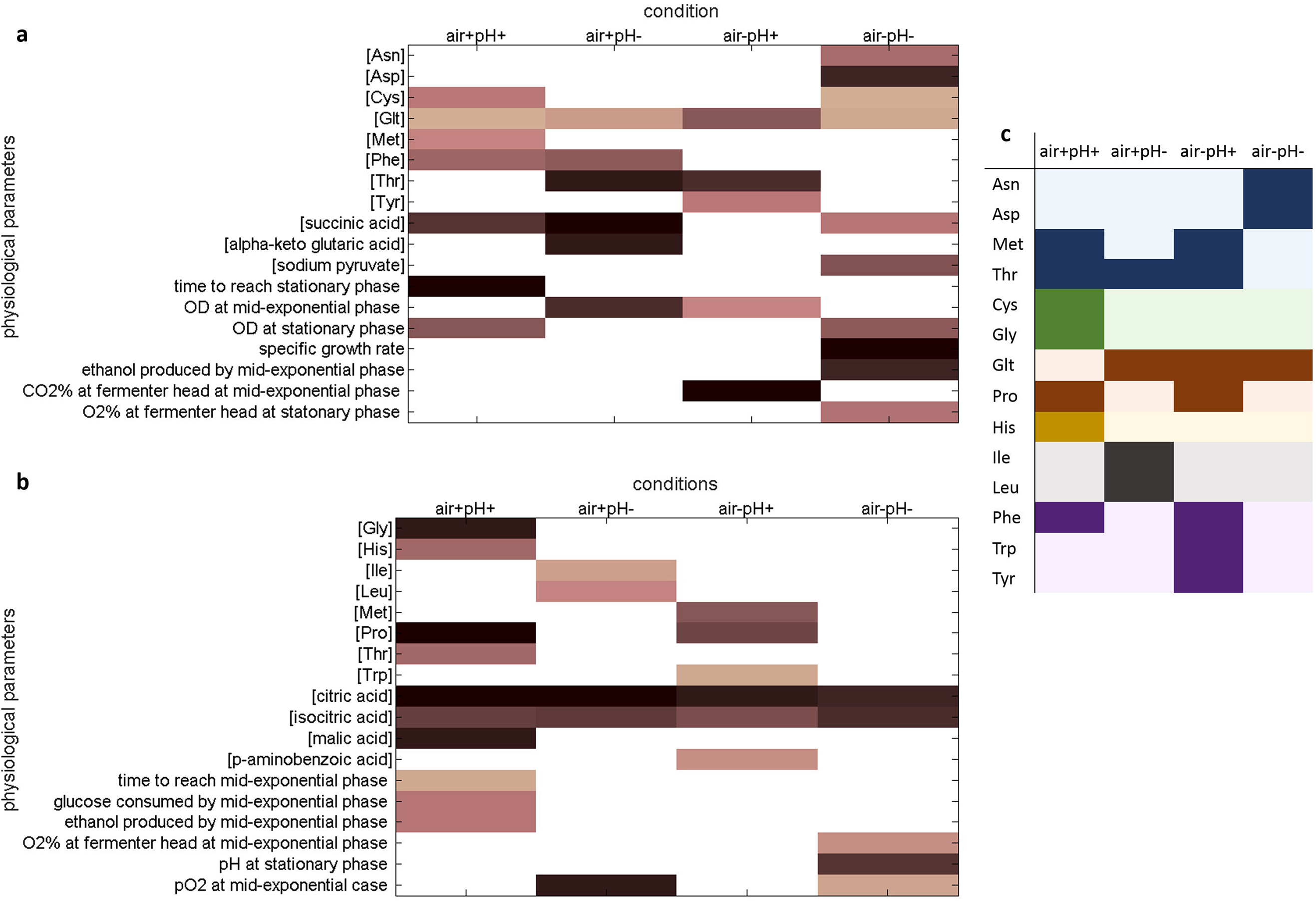
Culture condition-specific physiological response of yeast cells to long-term rapamycin exposure Physiological parameters that showed a significant difference in response to sustained rapamycin exposure in either one or more sets of culture conditions; air+pH+, air+pH-, air-pH+, and air-pH-, are displayed. (a) depicts the responses that become higher or faster in response to rapamycin exposure and (b) depicts those that are slowed down or lower than those for untreated controls. For colour coding in (a) and (b), see legend for Fig. 2. (c) denotes the amino acids, which display a significant change in the given culture conditions. Deeper shades denote a significant change and pale colours signify that there is no difference. The amino acids are colour-grouped into families that are derived from common molecules; shades of blue denote the aspartate family, shades of green denote the serine family, shades of red denote the glutamate family, shades of ochre denote histidine, shades of grey denote the pyruvate family, and the shades of purple denote the aromatic family.

The extracellular levels of amino acids derived from common molecules indicated a concerted action of the yeast amino acid metabolism when subjected to long-term rapamycin exposure. Either the members of a family were uniquely responsive under a specific set of conditions such as the serine, pyruvate, aromatic families, and histidine, or different members of the same family were responsive under all cultivation conditions tested, such as the aspartate and the glutamate families, which possess the additional role of generating precursors for amino acid biosynthesis (Fig. 3c).

### Transcriptional and endometabolic response of rapamycin-exposed yeast

Physiological response of yeast cells to long-term exposure to rapamycin designated differences in the extracellular availability of amino acid metabolites. Since the cells under investigation were only supplemented with histidine and leucine to address their auxotrophic requirements, the detection of amino acids in the culture supernatant was indicative of them as being produced and secreted as metabolites, potentially hinting towards global changes around the amino acid metabolism. We focused on investigating the transcriptional and endometabolic adaptation of the yeast cells to rapamycin, which led to the above discussed physiological differences (ESM3).

The global transcriptional arrangement of the yeast cells under the investigated conditions highlighted pH as the main factor in clustering of the gene expression profiles of different cultivations (Fig. 4a, b). A stricter similarity measure yielded tighter clusters, which classified the cultures according to the differences in their dissolved oxygen levels. The global transcriptional response of cells that have undergone long-term exposure to rapamycin was not substantially different from that of control cultures, indicative of the transcriptional adaptation of yeast cells (Fig 4c). Furthermore, the variation in the transcriptional response of those cultures inoculated into rapamycin-containing medium was lower than that of control cultures indicated by lower distance between replicate measurements. This observation was similar to the low variability observed for the preliminary cultures that were exposed to a similar mode of rapamycin administration (Fig. 1e).

**Fig.4.**
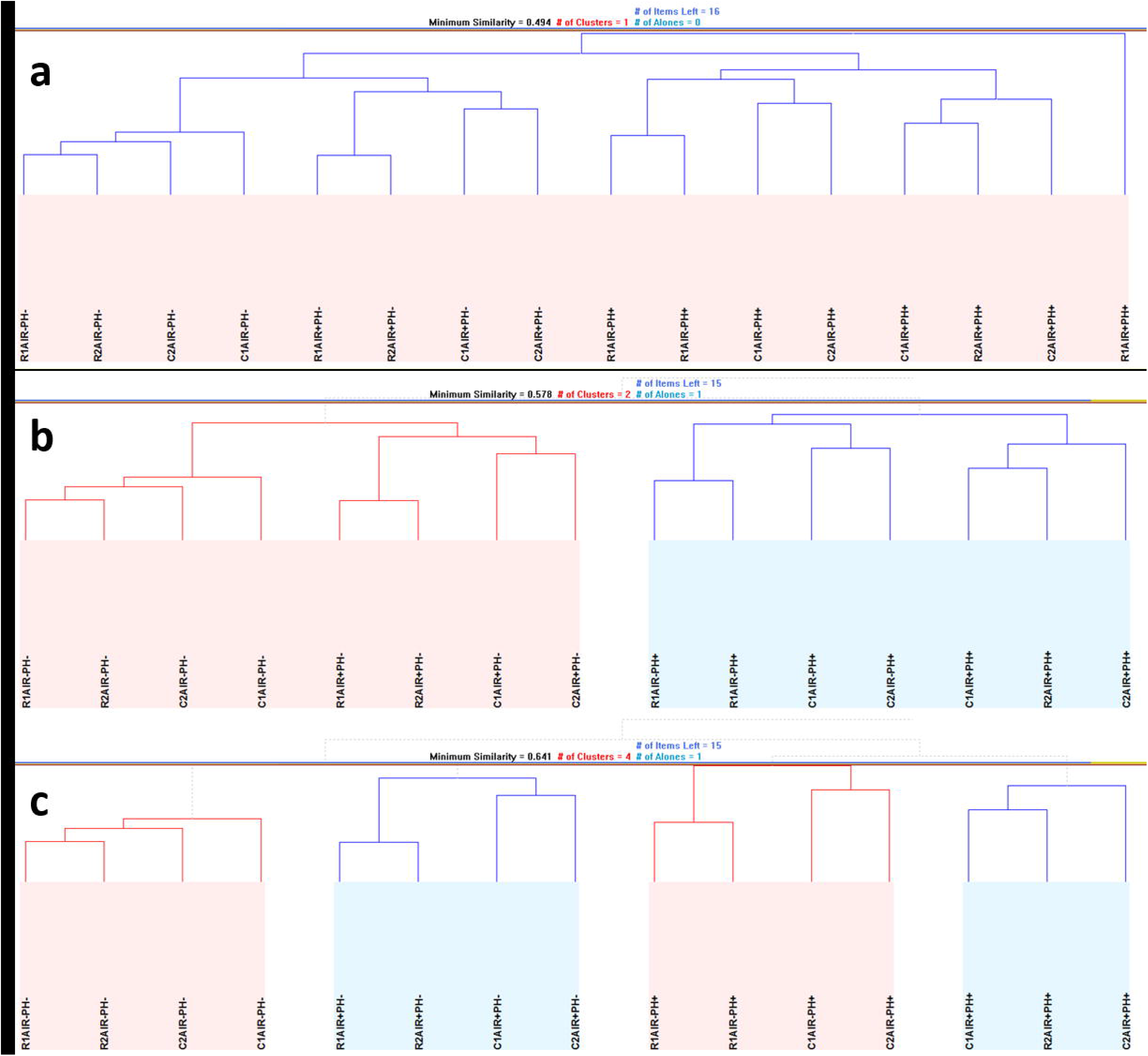
Hierarchical similarity of the global gene expression profiles among the cultures The hierarchical clustering of the transcriptome data for representing all 16 cultivations within a single cluster are shown at a minimum Pearson similarity distance of 0.494 (a). Two clusters denoted by pale blue and pale pink shaded areas are formed by employing a tighter metric than in (a) with a minimum Pearson similarity distance of 0.578. This measure leaves R1AIR+PH+ as a loner and the two clusters indicate differences in pH control among the cultures (b). Four clusters are formed of the remaining 15 cultures at an even tighter metric than that employed in (b) with a minimum Pearson similarity distance of 0.641, and these clusters are denoted by alternating pale blue and pale pink shaded areas. These clusters distinguish between the control of dissolved oxygen availability as well as pH (c). In all sub-figures, “C” denotes the control cultures and “R” denotes the rapamycin-treated cultures; “1” and “2” identify the replicates of the same experimental setup.

We investigated the transcriptional changes in response to rapamycin exposure in light of the environmental effectors; dissolved oxygen level, and extracellular pH. As a global response, genes whose expression levels significantly differed in magnitude (1.5-fold difference with significance threshold p-value < 5.00 × 10^−2^) were significantly enriched with ribonucleoprotein granule Cellular Component Gene Ontology term (p-value < 3.49 × 10^−2^). Among those genes, *SMX2* involved in mRNA splicing and capping, and *EAP1*, encoding an inhibitor of cap-dependent translation, also implicated in TOR signalling were upregulated, whereas *MET5* encoding a sulphite reductase involved in amino acid biosynthesis, *EBS1* involved in mRNA catabolic process, and *DHH1* suggested play a role in partitioning mRNAs between translatable and non-translatable pools were down-regulated. Furthermore, the gene expression of the signal recognition particle Srp102p, and Sec72p, involved in protein targeting and import into the endoplasmic reticulum were also elevated in cultures that experienced long-term exposure to rapamycin.

The genes that were downregulated (1.5-fold difference with significance threshold p-value < 5.00 × 10^−2^) in high dissolved oxygen availability (>90%) or low extracellular pH (<3.5) were significantly associated with cellular amino acid metabolic process Gene Ontology Term (p-value < 4.00 × 10^−2^). These genes were specifically involved with arginine and glutamine family amino acid metabolism, and particularly of biosynthetic routes, as well as related ornithine and urea metabolic processes [37]. This response was consistent with an environment-specific ANOVA analysis of rapamycin response at the transcriptional level; these pathways were significantly altered in air+pH− and air-pH- cultures (p-value < 1.00 × 10^−5^ for arginine and glutamine family pathways, and p-value < 1.00 × 10^−2^ for the urea cycle and ornithine metabolism). Since the role of arginine transport across the vacuolar membrane was implicated in the maintenance of acidic intracellular pH in *S. cerevisiae* with more than 90% of free arginine in the cell being located in the vacuole [38, 39], the downregulation of arginine biosynthesis could possibly be associated with the cells’ intracellular efforts to cope with low acidity extracellular environment.

Interestingly, exposure to rapamycin induced the gene expression of Mms21p and Pho81p, two proteins involved in protein serine/threonine kinase inhibitor activity. This functional enrichment was identified as significant among upregulated genes in cultures grown at relatively high extracellular pH (5.5) (p-value = 2.80 × 10^−4^). Although no direct physical interaction was reported for these proteins and the target of rapamycin serine/threonine kinase, *TOR1* gene was reported to be involved in an aggravating genetic interaction with *MMS21* (synthetic growth defect) [40]. The genes whose expression was significantly down-regulated in air-pH+ cultures were significantly enriched for reproductive process in single-celled organism Gene Ontology term (p-value = 4.86 × 10^−3^). Rapamycin was previously reported to block sexual development in fission yeast, so is a known repressor of reproductive processes through inhibition of a FKB12 homolog [41]. Although the *S. cerevisiae* homolog of this gene, *FPR2* [42] was differentially expressed in the presence of rapamycin (p-value < 5.0 × 10^−2^), the expression levels were very similar with and without treatment under the stated conditions. Furthermore, Mms21p was previously reported to prevent and eliminate dangerous recombination intermediates in meiosis in *S. cerevisiae* working together with Smc5/6p [43]. This antagonistic relationship between sexual reproduction routes and Mms21p was observed to be reinforced by prolonged exposure to rapamycin in cultures grown at pH = 5.5.

Genes, whose expression levels were significantly upregulated in response to rapamycin at or above 50% dissolved oxygen availability, and also at low extracellular pH (<3.5) were functionally associated with water transmembrane transporter activity (p-value < 2.00 × 10^−2^). Among these aquaporin-encoding genes, *AQY2* was reported to be selectively expressed in proliferating cells [44]. The relevance of aquaporins in yeast performance yet remains obscure [45], however, it has been shown that the selectivity of aquaporins depended highly on their constriction sites comprising a conserved arginine residue [46].

Concomitant with these changes in gene expression, endometabolic profiles of the cultures that were subjected to long-term rapamycin exposure were different from those for the control cultures. Intracellular amino acid and TCA intermediate concentrations displayed a coordinated response to sustained exposure to rapamycin. The intracellular succinate concentration of rapamycin treated cells was significantly higher than that of control similar to what has been observed in the extracellular environment of these cultures (p-value = 3.81 × 10^−4^). This was accompanied by lower malate levels indicating a draining of the TCA metabolic flow towards the end of the cycle (p-value = 6.63 × 10^−4^). Unlike the extracellular metabolome, intracellular citrate and isocitrate levels remained unchanged in response to long-term rapamycin-exposure. Significantly low α-ketoglutarate concentration in air+pH− cultures (p-value = 2.54 × 10^−3^) was coupled with high concentration of alanine, asparagine, arginine, glutamine and glutamate (p-value < 5.00 × 10^−2^), all produced from this TCA intermediate being converted first into glutamate. In contrast, high intracellular α-ketoglutarate concentration in rapamycin treated cultures (p-value < 1.30 × 10^−3^), was associated with either no accompanying change in glutamate and glutamate-family amino acid concentration (in air+pH+ cultures) or with low glutamate concentration (in air-pH- cultures) (p-value = 2.88 × 10^−3^).

## DISCUSSION

Rapamycin, is a potent inhibitor of the TOR kinase, which is the key nutrient-responsive controller of growth and aging in all eukaryotic cells. A pulse-like rapamycin treatment has long been known to exert a strong proteomic and transcriptional effect on eukaryotic cells from yeast [16, 23] to mammalian systems [47].

Furthermore, rapamycin was shown to extend lifespan and slow down aging in eukaryotic systems, although it has not been extensively investigated how the cells adapt themselves to prolonged exposure to the inhibitor. Here we investigated these effects at the phenotypical, metabolic and transcriptional levels, and associated this adaptation with the oxygen availability and the pH of the cells’ extracellular environment.

The changes observed in the amino acid profiles in response to oxygen levels was a response significantly enhanced for yeast cells, which were cultivated in the presence of rapamycin, whereas this effect was negligible for the untreated cells. Higher eukaryotes were shown to respond to oxygen availability more severely than yeast cells, which are able to sustain a healthy respiro-fermentative growth regime, do; rat heart cell metabolism was rewired away from protein biosynthesis towards the repletion of nucleotide pools by synthesis of purine nucleotides through the salvage pathway in response to lower oxygen availability [48]. Sustained rapamycin exposure possibly induced a response equivalent to elevated oxygen sensitivity in baker’s yeast, which is an inherently fitter organism to low oxygen availability than mammalian systems are. This response suggested the mimicking of the mammalian decision-making mechanisms around protein and nucleotide biosynthesis. A signal transduction pathway through mammalian TOR (mTOR) was identified previously to provide a checkpoint control, linking amino acid sufficiency to the control of peptide chain initiation in mammalian systems [49]. Here, we report a similar response induced by a repressor of the TOR pathway in microbial eukaryotes, with the major difference arising from the sourcing of the amino acids; metabolically synthesized instead of being imported.

Pronounced and significant differences were observed around the amino acid metabolism of yeast cells in response to rapamycin, and this response was frequently coupled with the environmental pH in which the cells were cultivated in. It was more prominent for the metabolism of amino acids, which yielded protein residues that were previously reported to be sensitive to pH levels, such as arginine [50]. mTOR Complex 1 (mTORC1) was previously shown to be involved in sensing lysosomal amino acids through a mechanism involving a vacuolar H^+^-ATPase [51]. The present findings suggest that the connection between pH and TOR could possibly be universal, and not only limited to mammalian systems. Role of free arginine and leucine in activation of the mTORC1 signalling pathway has been well-established [52], and the response of TOR signalling pathway to nutrient availability evaluated in light of this evidence [53]. Arginine and leucine are both essential amino acids for mammalian systems. Consequently the link between mTOR functionality and nutrient uptake plays a major role for these systems. The auxotrophy of the yeast cells employed here allowed the investigation of both essential and non-essential amino acid metabolic routes in this work. The arginine metabolism, particularly its biosynthetic pathway was shown to be affected in response to sustained repression of the TOR signalling pathway, and no significant metabolic changes were observed around the leucine metabolism, its uptake or transport, since this essential amino acid (for this particular strain) was available in abundance in the extracellular environment. These findings highlight some of the major differences in the nutrient-specific response of mammalian and microbial TOR functionality.

In conclusion, yeast cells were shown to adjust their physiology to sustained rapamycin exposure successfully and this adaptation was also accompanied by growth advantage. There were only a limited number of responses at the metabolic or at the transcriptional level as would be expected since extensive changes frequently encountered during rapamycin treatment would have been way too costly in terms of long-term cellular maintenance. However, these responses were persistently complementary across different levels of oxygen availability and extracellular acidity, implying a recurrent theme, particularly around the involvement of arginine metabolism in response to rapamycin. Another important aspect involved the decision making around smart resource allocation. Yeast cultures had to choose between inducing protein biosynthesis or nucleotide biosynthesis, and whether to spare the mRNAs to use in translation or to degrade them. All these subtle and minor but significant changes were responsible for the improvements in growth and longevity of the cultures that underwent sustained exposure to rapamaycin in the long run. Present findings on sustained exposure of yeast cells to rapamycin suggest the ability to synthesize amino acids and the decision making around resource allocation towards protein or nucleotide pool repletion constitute some of the major differences in cellular response of microbial eukaryotes and mammalian systems with dietary constraints around essential amino acids.

## ACKNOWLEDGEMENTS

Authors thank Ayca Cankorur Cetinkaya for critical reading of the manuscript.

Authors gratefully acknowledge the funding from the Leverhulme Trust (ECF-2016–681 to DD), TUBITAK (110M692 to BK), Turkish State Planning Organization (DPT-09K120520 to BK), and Bogazici University Research Fund (1932 to BK). None of the funding agencies had a role in study design, data collection or interpretation, or the decision to submit the work for publication. The authors declare no competing interests.

DD and ED designed the study and carried out the experimental work and DD conducted the statistical analyses and interpreted the results. SE carried out the metabolite and microarray analyses. DD, ED, and SE drafted the manuscript. BK and SGO conceived and coordinated the study, and helped draft the manuscript. All authors gave final approval for publication.

## ELECTRONIC SUPPLEMENTARY MATERIAL

**ESM1** Growth characteristics regarding the optimisation of the working concentration and the mode of administration of rapamycin (.xlsx format, single worksheet).

**ESM2** Growth characteristics and exometabolite levels of air±pH± cultures of yeast inoculated into 2nM rapamycin-containing medium (.xlsx format, seven worksheets including legend; data on fermentation characteristics, optical density, biomass density, exometabolite levels, exometabolite yields, and comparison of agitation at 400rpm vs 800rpm provided in separate worksheets).

**ESM3** Transcriptome and endometabolic profiles of air±pH± cultures of yeast inoculated into 2nM rapamycin-containing medium (.xlsx format, four worksheets including legend; gene expression profiles of rapamycin-treated and control cultures, the endometabolite profiles, and a summary of the significant Gene Ontology enrichments around the transcriptional changes identified provided in separate worksheets).

## References

1. Loewith R, Hall MN (2011) Target of rapamycin (TOR) in nutrient signaling and growth control. Genetics 189:1177–1201. doi: 10.1534/genetics.111.133363

2. Kapahi P, Chen D, Rogers AN, et al (2010) With TOR, less is more: a key role for the conserved nutrient-sensing TOR pathway in aging. Cell Metab 11:453–65. doi: 10.1016/j.cmet.2010.05.001

3. Castrillo JI, Zeef LA, Hoyle DC, et al (2007) Growth control of the eukaryote cell: a systems biology study in yeast. J Biol 6:4. doi: 10.1186/jbiol54

4. Loewith R, Hall MN (2011) Target of rapamycin (TOR) in nutrient signaling and growth control. Genetics 189:1177–201. doi: 10.1534/genetics.111.133363

5. Lamming DW, Ye L, Sabatini DM, Baur JA (2013) Rapalogs and mTOR inhibitors as anti-aging therapeutics. J Clin Invest 123:980–989. doi: 10.1172/JCI64099

6. Li J, Kim SG, Blenis J (2014) Rapamycin: one drug, many effects. Cell Metab 19:373–9. doi:10.1016/j.cmet.2014.01.001

7. Barbet NC, Schneider U, Helliwell SB, et al (1996) TOR controls translation initiation and early G1 progression in yeast. Mol Biol Cell 7:25–42

8. Huber A, Bodenmiller B, Uotila A, et al (2009) Characterization of the rapamycin-sensitive phosphoproteome reveals that Sch9 is a central coordinator of protein synthesis. Genes Dev 23:1929–1943. doi: 10.1101/gad.532109

9. Cardenas ME, Cutler NS, Lorenz MC, et al (1999) The TOR signaling cascade regulates gene expression in response to nutrients. Genes Dev 13:3271–9

10. Conrad M, Schothorst J, Kankipati HN, et al (2014) Nutrient sensing and signaling in the yeast *Saccharomyces cerevisiae*. FEMS Microbiol Rev 38:254–299. doi: 10.1111/1574-6976.12065

11. Powers RW, Kaeberlein M, Caldwell SD, et al (2006) Extension of chronological life span in yeast by decreased TOR pathway signaling. Genes Dev 20:174–184. doi: 10.1101/gad.1381406.A

12. Anisimov VN, Zabezhinski MA, Popovich IG, et al (2010) Rapamycin extends maximal lifespan in cancer-prone mice. Am J Pathol 176:2092–7. doi: 10.2353/ajpath.2010.091050

13. Anisimov VN, Zabezhinski MA, Popovich IG, et al (2011) Rapamycin increases lifespan and inhibits spontaneous tumorigenesis in inbred female mice. Cell Cycle 10:4230–6. doi: 10.4161/cc.10.24.18486

14. Popovich IG, Anisimov VN, Zabezhinski MA, et al (2014) Lifespan extension and cancer prevention in HER-2/neu transgenic mice treated with low intermittent doses of rapamycin. Cancer Biol Ther 15:586–92. doi: 10.4161/cbt.28164

15. Chen L, Madura K (2014) Degradation of specific nuclear proteins occurs in the cytoplasm in Saccharomyces cerevisiae. Genetics 197:193–7. doi: 10.1534/genetics.114.163824

16. Hardwick JS, Kuruvilla FG, Tong JK, et al (1999) Rapamycin-modulated transcription defines the subset of nutrient-sensitive signaling pathways directly controlled by the Tor proteins. Proc Natl Acad Sci 96:14866–14870. doi: 10.1073/pnas.96.26.14866

17. Zhang N, Quan Z, Rash B, Oliver SG (2013) Synergistic effects of TOR and proteasome pathways on the yeast transcriptome and cell growth. Open Biol 3:120137–120137. doi: 10.1098/rsob.120137

18. Hughes Hallett JE, Luo X, Capaldi AP (2014) State transitions in the TORC1 signaling pathway and information processing in Saccharomyces cerevisiae. Genetics 198:773–86. doi:10.1534/genetics.114.168369

19. Bolstad BM, Irizarry RA, Astrand M, Speed TP (2003) A comparison of normalization methods for high density oligonucleotide array data based on variance and bias. Bioinformatics 19:185–193. doi: 10.1093/bioinformatics/19.2.185

20. Reinke A, Chen JC-Y, Aronova S, Powers T (2006) Caffeine targets TOR complex I and provides evidence for a regulatory link between the FRB and kinase domains of Tor1p. J Biol Chem 281:31616–31626. doi: 10.1074/jbc.M603107200

21. Rallis C, Codlin S, Bähler J (2013) TORC1 signaling inhibition by rapamycin and caffeine affect lifespan, global gene expression, and cell proliferation of fission yeast. Aging Cell 12:563–73. doi:10.1111/acel.12080

22. Yeung KY, Dombek KM, Lo K, et al (2011) Construction of regulatory networks using expression time-series data of a genotyped population. Proc Natl Acad Sci U S A 108:19436–41. doi:10.1073/pnas.1116442108

23. Oliveira AP, Ludwig C, Zampieri M, et al (2015) Dynamic phosphoproteomics reveals TORC1-dependent regulation of yeast nucleotide and amino acid biosynthesis. Sci Signal 8:rs4. doi:10.1126/scisignal.2005768

24. Evans SK, Burgess KE V., Gray J V. (2014) Recovery from Rapamycin. J Biol Chem 289:26554–26565. doi: 10.1074/jbc.M114.589754

25. Dai X, Yan H, Li N, et al (2016) Metabolic adaptation of microbial communities to ammonium stress in a high solid anaerobic digester with dewatered sludge. Sci Rep 6:28193. doi: 10.1038/srep28193

26. Wang X, Kang Y, Luo C, et al (2014) Heteroresistance at the single-cell level: adapting to antibiotic stress through a population-based strategy and growth-controlled interphenotypic coordination. MBio 5:e00942–13. doi: 10.1128/mBio.00942-13

27. Torsvik V, Øvreås L (2008) Microbial Diversity, Life Strategies, and Adaptation to Life in Extreme Soils. Springer, Berlin, Heidelberg, pp 15–43

28. Bell G (2013) Evolutionary rescue and the limits of adaptation. Philos Trans R Soc Lond B Biol Sci 368:20120080. doi: 10.1098/rstb.2012.0080

29. Ehninger D, Neff F, Xie K (2014) Longevity, aging and rapamycin. Cell Mol Life Sci 71:4325–46. doi:10.1007/s00018-014-1677-1

30. Neff F, Flores-Dominguez D, Ryan DP, et al (2013) Rapamycin extends murine lifespan but has limited effects on aging. J Clin Invest 123:3272–91. doi: 10.1172/JCI67674

31. Madrid M, Vázquez-Marín B, Franco A, et al (2016) Multiple crosstalk between TOR and the cell integrity MAPK signaling pathway in fission yeast. Sci Rep 6:37515. doi: 10.1038/srep37515

32. Rintala E, Toivari M, Pitkänen J-P, et al (2009) Low oxygen levels as a trigger for enhancement of respiratory metabolism in Saccharomyces cerevisiae. BMC Genomics 10:461. doi: 10.1186/1471-2164-10-461

33. Burtner CR, Murakami CJ, Kennedy BK, Kaeberlein M (2009) A molecular mechanism of chronological aging in yeast. Cell Cycle 8:1256–1270

34. Bonawitz ND, Chatenay-Lapointe M, Pan Y, Shadel GS (2007) Reduced TOR Signaling Extends Chronological Life Span via Increased Respiration and Upregulation of Mitochondrial Gene Expression. Cell Metab 5:265–277. doi: 10.1016/j.cmet.2007.02.009

35. Murakami CJ, Wall V, Basisty N, Kaeberlein M (2011) Composition and acidification of the culture medium influences chronological aging similarly in vineyard and laboratory yeast. PLoS One 6:e24530–e24530. doi: 10.1371/journal.pone.0024530

36. Thomas KC, Hynes SH, Ingledew WM (2002) Influence of medium buffering capacity on inhibition of Saccharomyces cerevisiae growth by acetic and lactic acids. Appl Environ Microbiol 68:1616–1623

37. Ljungdahl PO, Daignan-Fornier B (2012) Regulation of amino acid, nucleotide, and phosphate metabolism in Saccharomyces cerevisiae. Genetics 190:885–929. doi: 10.1534/genetics.111.133306

38. Cooper SJ, Finney GL, Brown SL, et al (2010) High-throughput profiling of amino acids in strains of the Saccharomyces cerevisiae deletion collection. Genome Res 20:1288–96. doi:10.1101/gr.105825.110

39. Wykoff DD, O’Shea EK, Daignan-Fornier B, André B (2001) Phosphate transport and sensing in Saccharomyces cerevisiae. Genetics 159:1491–9. doi: 10.1534/genetics.166.4.1727

40. Alam MT, Merlo ME, STREAM Consortium TSC, et al (2010) Metabolic modeling and analysis of the metabolic switch in Streptomyces coelicolor. BMC Genomics 11:202. doi: 10.1186/1471-2164-11-202

41. Weisman R, Finkelstein S, Choder M (2001) Rapamycin blocks sexual development in fission yeast through inhibition of the cellular function of an FKBP12 homolog. J Biol Chem 276:24736–42. doi:10.1074/jbc.M102090200

42. Partaledis JA, Fleming MA, Harding MW, Berlin V (1992) Saccharomyces cerevisiae contains a homolog of human fkbp-13, a membrane-associated fk506/rapamycin binding protein. Yeast 8:673–680. doi: 10.1002/yea.320080812

43. Xaver M, Huang L, Chen D, Klein F (2013) Smc5/6-Mms21 Prevents and Eliminates Inappropriate Recombination Intermediates in Meiosis. PLoS Genet 9:e1004067. doi: 10.1371/journal.pgen.1004067

44. Pettersson N, Filipsson C, Becit E, et al (2005) Aquaporins in yeasts and filamentous fungi. Biol Cell 97:487–500. doi: 10.1042/BC20040144

45. Soveral G, Prista C, Moura TF, Loureiro-Dias MC (2011) Yeast water channels: an overview of orthodox aquaporins. Biol Cell 103:35–54. doi: 10.1042/BC20100102

46. Kruse E, Uehlein N, Kaldenhoff R (2006) The aquaporins. Genome Biol 7:206. doi: 10.1186/gb-2006-7-2-206

47. Hukelmann JL, Anderson KE, Sinclair L V, et al (2016) The cytotoxic T cell proteome and its shaping by the kinase mTOR. Nat Immunol 17:104–12. doi: 10.1038/ni.3314

48. Ravid K, Diamant P, Avi-Dor Y (1985) Interrelation between salvage of purine nucleotides and protein synthesis in rat heart cells. Arch Biochem Biophys 236:159–166. doi: 10.1016/0003-9861(85)90615-0

49. Hara K, Yonezawa K, Weng QP, et al (1998) Amino acid sufficiency and mTOR regulate p70 S6 kinase and eIF-4E BP1 through a common effector mechanism. J Biol Chem 273:14484–94

50. White KA, Ruiz DG, Szpiech ZA, et al (2017) Cancer-associated arginine-to-histidine mutations confer a gain in pH sensing to mutant proteins. Sci Signal 10:eaam9931. doi: 10.1126/scisignal.aam9931

51. Zoncu R, Bar-Peled L, Efeyan A, et al (2011) mTORC1 Senses Lysosomal Amino Acids Through an Inside-Out Mechanism That Requires the Vacuolar H+-ATPase. Science (80-) 334:678–683. doi:10.1126/science.1207056

52. Wyant GA, Abu-Remaileh M, Wolfson RL, et al (2017) mTORC1 Activator SLC38A9 Is Required to Efflux Essential Amino Acids from Lysosomes and Use Protein as a Nutrient. Cell 171:642–654.e12. doi: 10.1016/j.cell.2017.09.046

53. Gu X, Orozco JM, Saxton RA, et al (2017) SAMTOR is an *S* –adenosylmethionine sensor for the mTORC1 pathway. Science (80-) 358:813–818. doi: 10.1126/science.aao3265

54. Brachmann CB, Davies A, Cost GJ, et al (1998) Designer deletion strains derived from Saccharomyces cerevisiae S288C: a useful set of strains and plasmids for PCR-mediated gene disruption and other applications. Yeast 14:115–32. doi: 10.1002/(SICI)1097-0061(19980130)14:2<115::AIDYEA204>3.0.CO;2-2

55. Baganz F, Hayes A, Marren D, et al (1997) Suitability of replacement markers for functional analysis studies in Saccharomyces cerevisiae. Yeast 13:1563–73. doi: 10.1002/(SICI)1097-0061(199712)13:16<1563::AID-YEA240>3.0.CO;2-6

56. Fabrizio P, Longo VD (2007) The Chronological Life Span of Saccharomyces cerevisiae. Humana Press, pp 89–95

57. Li C, Wong WH (2001) Model-based analysis of oligonucleotide arrays: expression index computation and outlier detection. Proc Natl Acad Sci U S A 98:31–36. doi: 10.1073/pnas.011404098

58. Brazma A, Hingamp P, Quackenbush J, et al (2001) Minimum information about a microarray experiment (MIAME)-toward standards for microarray data. Nat Genet 29:365–371. doi:10.1038/ng1201-365

59. Villas-Bôas SG, Højer-Pedersen J, Akesson M, et al (2005) Global metabolite analysis of yeast: evaluation of sample preparation methods. Yeast 22:1155–1169. doi: 10.1002/yea.1308

60. Seo J, Bakay M, Chen Y-W, et al (2004) Interactively optimizing signal-to-noise ratios in expression profiling: project-specific algorithm selection and detection p-value weighting in Affymetrix microarrays. Bioinformatics 20:2534–44. doi: 10.1093/bioinformatics/bth280

61. Boyle EI, Weng S, Gollub J, et al (2004) GO::TermFinder-open source software for accessing Gene Ontology information and finding significantly enriched Gene Ontology terms associated with a list of genes. Bioinformatics 20:3710–5. doi: 10.1093/bioinformatics/bth456

